# Complex extracellular biology drives surface competition in *Bacillus subtilis*

**DOI:** 10.1101/2022.02.28.482363

**Authors:** Theresa Jautzus, Jordi van Gestel, Ákos T. Kovács

**Author notes:** These authors contributed equally to this work (Theresa Jautzus, Jordi van Gestel).

## Abstract

In nature, many bacteria grow on surfaces, where they form cell collectives that compete for space. Within these collectives, cells often secrete molecules that benefit surface spreading by, for example, reducing surface tension or promoting filamentous growth. Although we have a detailed understanding of how these molecules are produced, much remains unknown about their role in surface competition. Here, we examine sliding motility in *Bacillus subtilis* and compare how secreted molecules, essential for sliding, affect cooperation and competition on the surface. We specifically examine (i) the lipopeptide surfactin, (ii) the hydrophobin protein BslA and (iii) exopolysaccharides (EPS). We find that these molecules have a remarkably different effect on competition: whereas surfactin acts like a common good, which is costly to produce and benefits cells throughout the surface, BslA and EPS are cost-free and act locally. Accordingly, surfactin deficient mutants can exploit the wild-type strain in competition for space, while BslA and EPS mutants cannot. Using a mathematical model, we show that three factors are important in predicting the outcome of surface competition: the costs of molecule synthesis, the private benefits of molecule production, and the diffusion rate. Our results underscore the intricate extracellular biology that can drive bacterial surface competition.

## Introduction

Surface-associated bacteria can be found throughout ecology, from the cells adhering to our teeth to the dense bacterial colonies growing on plant roots. On the surface, cells form collectives that strongly compete for space. In doing so, cells often secrete antimicrobial compounds that antagonize competitors on the surface (e.g. [1]). Cells can also work together and thereby promote collective growth or dispersal [2–6]. This is perhaps best illustrated when it comes to surface expansion. In many bacterial species, cells move in a coordinated fashion. Depending on the driving forces, this form of collective movement is referred to as swarming, gliding, or sliding [7]. For example, in the case of swarming, cells collectively express flagella that advance the colony boundary through propulsion. Similarly, bacteria can also slide over surfaces, which is driven by cell division and occurs independently from cellular appendages like flagella. For example, in *Bacillus subtilis*, sliding motility results from cells forming filamentous bundles that push themselves away from the colony center as they grow [8].

Many forms of collective movement, including swarming and sliding, dependent on the secretion of extracellular molecules [9–12]. For example, in the case of *B. subtilis*, sliding depends on the production and secretion of the lipopeptide surfactin, the bacterial hydrophobin protein BslA, and exopolysaccharides (EPS) [13, 14]. Without these molecules, colonies cannot spread. The secreted molecules carryout different functions: surfactin for example acts as a surfactant and thereby lowers the surface tension, which facilitates lateral expansion. Through its hydrophobic properties [15, 16], BslA is assumed to have a similar biophysical effect. In contrast, EPS is required for the formation and expansion of filamentous bundles [8]. EPS might furthermore promote colony expansion by increasing the osmotic pressure within the colony [17].

Despite promoting surface expansion, cells that produce the above molecules do not necessarily gain a competitive advantage. For instance, when molecules are costly to produce and benefit other cells on the surface, producers might be exploited by close-by non-producers. This results in a so-called common goods problem, where the production of a common good, in this case a secreted molecule that promotes surface expansion, is selected against despite its beneficial effect on surface spreading. For example, in the case of *B. subtilis* colony biofilm growth, it was previously shown that EPS production promotes expansion, but is nevertheless selected against because EPS-deficient cells outgrow EPS-producing cells in competition for space [6].

There are several factors that can make it challenging to determine whether secreted molecules act as common goods. First, metabolic costs of molecule synthesis can differ between conditions. When molecule production is only costly under specific conditions (i.e. nutrient limitation), there is only a risk of exploitation under those conditions [18–20]. Second, the spatial scale at which molecules are shared may vary [21]. For instance, *B. subtilis* cells broadly share EPS molecules during both complex colony development and pellicle formation [22–30], however, when it comes to sliding, EPS deficient mutants are excluded from the expanding colony edge [8], which suggests that EPS might not be broadly shared during sliding conditions. Cells can also directly limit the diffusion of secreted molecules themselves. For example, Drescher and colleagues [5] demonstrated that *Vibrio cholerae* cells limit the diffusion of chitinase by forming dense biofilms. In the absence of biofilm formation, chitinase production, which benefits surface growth by degrading chitin, is exploited by non-producing cells. Also, it was recently shown that the diffusion of quorum-sensing signals can be limited by active signal uptake [31]. Finally, secreted molecules may also provide directly benefits to the producer, thereby functioning as private or semi-common goods. For instance, in the budding yeast *Saccharomyces cerevisiae*, cells secrete an enzyme called invertase to hydrolyze sucrose. Although most monosaccharides produced by hydrolysis diffuse away, about 1% is immediately taken up by cell, thereby providing an immediate private benefit [32].

The above examples illustrate that the extracellular biology of surface growth can impact competition in versatile ways. In this study, we therefore investigate how the secreted molecules, essential for sliding motility in *B. subtilis*, affect spatial competition. We performed repeated competition experiments, using both knockout strains and inducible strains that differ in the production and secretion of extracellular molecules. We subsequently used a spatial-explicit model, based on minimal number of mechanistic assumptions, to recapitulate the experimental results. Together, the experiments and model reveal that three factors are key in determining the outcome of competition: the costs of molecule synthesis, the private benefits of molecule production and the diffusion rate.

## Materials and Methods

### Strains and cultivating conditions

All *B. subtilis* strains used in this study are derivatives of a competence enhanced NCBI3610 (DK1042 [33]). A Δ*hag* mutant lacking flagellin was used as “sliding wild type” since motile strains would swarm and sliding could not be investigated under the used conditions. The strains used in this study are listed in Table S1. To engineer mutant derivatives, a *B. subtilis* receptor strain was transformed with genomic DNA from a donor strain (method after [33]) or with the respective plasmid. A detailed description of the strain construction can be found in the Supplemental material, used primers listed in Table S2. The following antibiotic concentrations were used for respective resistant strains if appropriate: ampicillin 100 µg/ml (Amp), spectinomycin 100 µg/ml (Spec), kanamycin 5 µg/ml (Km), chloramphenicol 5 µg/ml (Cm), tetracycline 10 µg/ml (Tet), MLS: erythromycin 1 µg/ml + lincomycin 12.5 µg/ml.

### Sliding competition assay

Sliding assays were performed as described previously [34]. Briefly, overnight cultures of *B. subtilis* were density normalized and if required, mixed in a 1:1, 1:10 or 10:1 ratio with a competitor strain. 2 µl of the strain or mixture were spotted on semi-solid lysogeny broth (LB, Lennox, Carl Roth) supplemented with 0.7 % agar in 9 cm diameter plates that were dried 20 min prior and 10 min post inoculation. The plates were incubated at 37°C for 24 h if not stated otherwise and sliding was evaluated by assessing the diameter of the expanded colony. Additionally, distribution of fluorescently labeled strains within the sliding disk was evaluated by detecting the fluorescence signal with an AxioZoom V16 fluorescence stereomicroscope equipped with a Zeiss CL 9000 LED light source, HE eGFP filter set (excitation at 470/40 nm and emission at 525/50 nm), HE mRFP filter set (excitation at 572/25 nm and emission at 629/62 nm) and an AxioCam MRm monochrome camera (Carl Zeiss Microscopy GmbH; for exact details of the instrument, see [28]). For image display, the brightness of all fluorescence images was adjusted in the same way and the background was subtracted using the program ImageJ with the rolling ball option (1100 pixels radius).

### Determination of occupied area in the sliding colony

The area each strain occupies in the sliding colony was determined using fluorescence stereomicroscope images of the mixtures of different fluorescently labelled strains and the software ImageJ (detailed description in [34]). Briefly, the images were opened in ImageJ, separated by channel and the scale was changed to pixel. After background removal, a defined threshold was applied to the image of the green and red channel to separate the fluorescence signal corresponding to the occupied area of the respective strain. The total area above the threshold was selected and measured as number of pixels. To compare both strains in the sliding colony, the ratio of the two areas was calculated. A summary of the area and percentage calculation was not possible since there was often at least a small overlap between the areas of the different strains, especially in the center of the sliding colony. For this analysis, original images were used.

### Complementation assay

To complement sliding of mutants with externally supplied goods, cultures and plates were prepared as described above (see Sliding competition assay). Additionally, 2 µl of the respective compound was spotted 5 min before the mutant culture (TB532, TB534, TB536) on the same inoculation point. The respective mutant and wild-type cultures alone were used as controls. For surfactin complementation, 10, 1, 0.1 and 0.01 mg/ml of commercial surfactin (Sigma Aldrich) dissolved in methanol was used and pure methanol was spotted as control.

For BslA complementation, the lysate of a BslA-producing *Escherichia coli* BL21 strain (NRS4110 containing plasmid pNW1128 [16]) and an *E. coli* BL21 strain without the BslA-production plasmid was used. To obtain the *E. coli* lysate, the strains were grown overnight in LB medium and were afterwards inoculated 1:100 in autoinduction medium [35]. The NRS4110 culture was always supplemented with 100 µg/ml Amp. The cultures were grown for 7-8 h at 37°C with 225 rpm shaking and cells were collected by centrifugation at 4000 g for 15 min. The pellet was resuspended in 5ml PBS buffer supplemented with 1 mM EDTA and cooled on ice for 10 min. The suspension was then sonicated using an Ultrasonic Processor VCX-130 (Zinsser Analytics, Frankfurt am Main, Germany) with ten repeats of a 10 s pulse of 45 % amplitude. During sonication, the suspension was cooled on ice. To obtain the lysate, the suspension was centrifuged (5000 g, 15 min), the supernatant was collected, filter sterilized and stored at 4°C.

For exopolysaccharide complementation, EPS was isolated from pellicle biofilms of a *B. subtilis* NCIB 3610 wild-type strain, a Δ*tasA* and as control a Δ*eps* mutant with slight modifications after [36]. Briefly, four wells of a 24-well plate containing 2 ml biofilm promoting liquid MSgg medium each [37] were inoculated 1:100 with overnight culture of the respective strain and incubated at 30°C for 2-3 d. The pellicle formed at the air-liquid interface was collected together with the medium, diluted 1:1 with PBS buffer and vortexed. The pellicles were sonicated (Ultrasonic Processor VCX-130, Zinsser Analytics, Frankfurt am Main, Germany; 2 × 12 pulses of 1 s with 30% amplitude), 0.2 M NaOH was added to a final concentration of 0.1 M and the samples were incubated at room temperature for 10 min with short periodic vortexing. Afterwards, the samples were chilled on ice for 5 min before adding cold 0.4 M HCl to a final concentration of 0.1 M. The samples were centrifuged (7000 g for 15 min at 4°C), the supernatant was collected and transferred to a three-fold volume of cold 96% ethanol and incubated for ca. 20 h at 4°C. The precipitated EPS was collected by centrifugation (7000 g for 15 min at 4°C) and the pellet was dried overnight at 55°C. After drying, the EPS was re-dissolved with deionized water in a 1:10 ratio and NaCl was added to a final concentration of 0.5%. The precipitation, collection and dissolving step was repeated once and the isolated EPS was filter sterilized and stored at 4°C. The functionality of the BslA-containing lysate and the isolated EPS was successfully verified by testing them for biofilm formation.

### Fitness assay

To determine the relative fitness under sliding conditions, strains with inducible gene constructs of *epsA, bslA* and *srfAA* (TB875, TB873, TB977, respectively) were competed against the respective mutants (TB893, TB922, TB895, respectively). Therefore, the strains were density normalized and mixed in a 1:1 ratio in a reaction tube. This mixture was used to a) determine the colony forming units (cfu) at the start by plating on antibiotic containing plates selective for each strain and b) to inoculate a 50 ml Schott bottle containing 5 ml LB medium in a 1:100 dilution. The cultures were incubated at 37°C and 225 rpm shaking for 6-6 ½ h after which they were diluted 1:100 and grown again for 6-6 ½ h under the same conditions. Following incubation, the final cfu of the two competing strains was determined as described above. Initial and final cfu were then used to calculate the relative fitness via the so called malthusian parameter after Lenski *et a*l. [38] with r = (m inducible strain)/(m mutant) and m = ln [(final cfu)/(initial cfu)].

### Mathematical model

To further investigate surface competition, we construed an agent-based model to simulate surface growth with the goal of recapitulating the experimental results and thereby identifying the factors that shape the outcome of competition. In brief, we modeled a two-dimensional surface on which cells can grow. The surface consists of a hexagonal grid of 100×100 elements, which together are 100mm in width, approximating the diameter of a Petri dish. Like for our experiments, at the start of growth, we assume resources are homogeneously distributed (*R*_*init*_) and cells occur at the surface center (inoculum is 10mm in diameter). Cells consume resources (*R*) and divide at a constant rate: following our experiments, wild-type cells divide approximately every 45 minutes (*r*_*wt*_) and *srf* mutants have a slight ∼3% fitness advantage (*r*_*srf*_). Cell division is impossible when resources are depleted (*R* < 1). As observed in our experiments, a colony can only expand in the presence of surfactin (*M*_*1*_), EPS (*M*_*2*_) *and* BslA (*M*_*3*_) production. We assume that the wild type secretes all these molecules at a constant rate (*d*), while the knockout mutants (Δ*srfAA*, Δ*eps*, Δ*bslA*) fail to produce one of these respective molecules. The secreted molecules degrade at a constant rate (δ) and diffuse in space (α). Resembling sliding motility, we assume that if the local concentration of surfactin, EPS *and* BslA are sufficiently high (*M*_*1*_, *M*_*2*_, *M*_*3*_ ≥ ⊤, threshold) a daughter cells is placed in neighboring grid element upon cell division, leading to lateral colony expansion. If the concentration of any of the three molecules is too low (min(*M*_*1*_,*M*_*2*_,*M*_*3*_) < ⊤), daughter cells remain on the same grid element as the mother cell and the colony is thought to expand vertically. We implement any private benefits that may result from molecule production as a reduced concentration at which colony expansion is possible. For example, when wild-type cells have a 20% private benefit from secreting EPS, we assume these cells expand at a 20% lower EPS concentration compared to *eps*-mutant cells – thereby having a direct competitive advantage, despite having the same of cell division rate (note that we did not observe differences in cell division rates between wild type, Δ*eps* and Δ*bslA* mutants).

Since surfactin functions as a classical common good, we use our model to study the more-intricate functional implications of EPS and BslA production. We first explore parameter space by varying the diffusion rates of EPS and BslA (10^−8^, 10^−6^, 10^−4^, 10^−2^ mm^2^ s^-1^) as well as their private benefits (0, 10, 50, 90%). For each parameter condition, we simulate seven pairwise competitions: (1) WT-WT, (2) WT-Δ*eps*, (3) WT-Δ*bslA*, (4) WT-Δ*srfAA*, (5) Δ*eps-*Δ*bslA*, (6) Δ*eps-*Δ*srf*, (7) Δ*bslA-*Δ*srf*. For each simulation, we started with an equal number of cells from each genotype, let them grow for 24 hours, and then quantified their abundance in the colony as well as the colony size. Second, to determine how well the simulations reproduced our experimental results, we scored the overall match between our simulation results and experimental results: (1) WT-WT: strains should be equally abundant at end of competition; (2) WT-Δ*srf*: Δ*srf* mutant wins competition; (3) WT-Δ*eps*: wild type wins competition; (4) WT-Δ*bslA*: wild type wins competition, but colonies are slightly smaller than those of the wild type; (5) Δ*eps*-Δ*bslA*: Δ*bslA* mutant wins competition and produce the smallest observed colonies; (6) Δ*eps*-Δ*srf*: Δ*srf* mutant wins competition: (7) Δ*bslA*-Δ*srf*: Δ*srf* mutant wins competition. For each parameter conditions, we assign a score from 0 to 7, where 0 means that none of the competition outcomes were correctly predicted and 7 means that they were all correctly predicted. We determined the scores across parameter space using four replicate simulations per condition. For the simulations that best matched our experimental results, we also studied the distribution of surfactin, EPS and BslA at the end of colony growth. In the supplementary data, we provide the lists of both parameters (Table S3) and variables (Table S4); as well as a pseudocode description of our simulations. The C++ code for running the simulations is available on GitHub: https://github.com/jordivangestel/Bacillus-subtilis-surface-competition.

## Results

### Extracellular molecules have diverse effects on spatial competition

Sliding motility is driven by cell division, thereby occurring independently from flagella propulsion. During sliding, the colony grows as a flat circular disk that expands outwards over a surface. To assure that cells express sliding motility, without swimming or swarming, we exclusively examined *B. subtilis* strains that are deficient in flagellin production (Δ*hag*). Thus, whenever we refer to a wild-type or mutant strain, this strain contains a *hag* deletion.

We started our analysis by examining the wild-type and non-producing mutant strains in isolation, by growing them for 24 h on a semi-solid surface. We consider mutants deficient in EPS, BslA or surfactin (Δ*eps*, Δ*bslA*, Δ*srfAA*, respectively) production. After a short lag-phase, the wild-type colony slid over the surface at a constant rate, reaching a diameter of about 3 cm after 24 h (Fig. 1; see also [34]). In contrast, and in agreement with previous studies [14], the non-producing mutants were strongly impaired in colony spreading, reaching a diameter of ∼1 cm only.

**Fig. 1.**
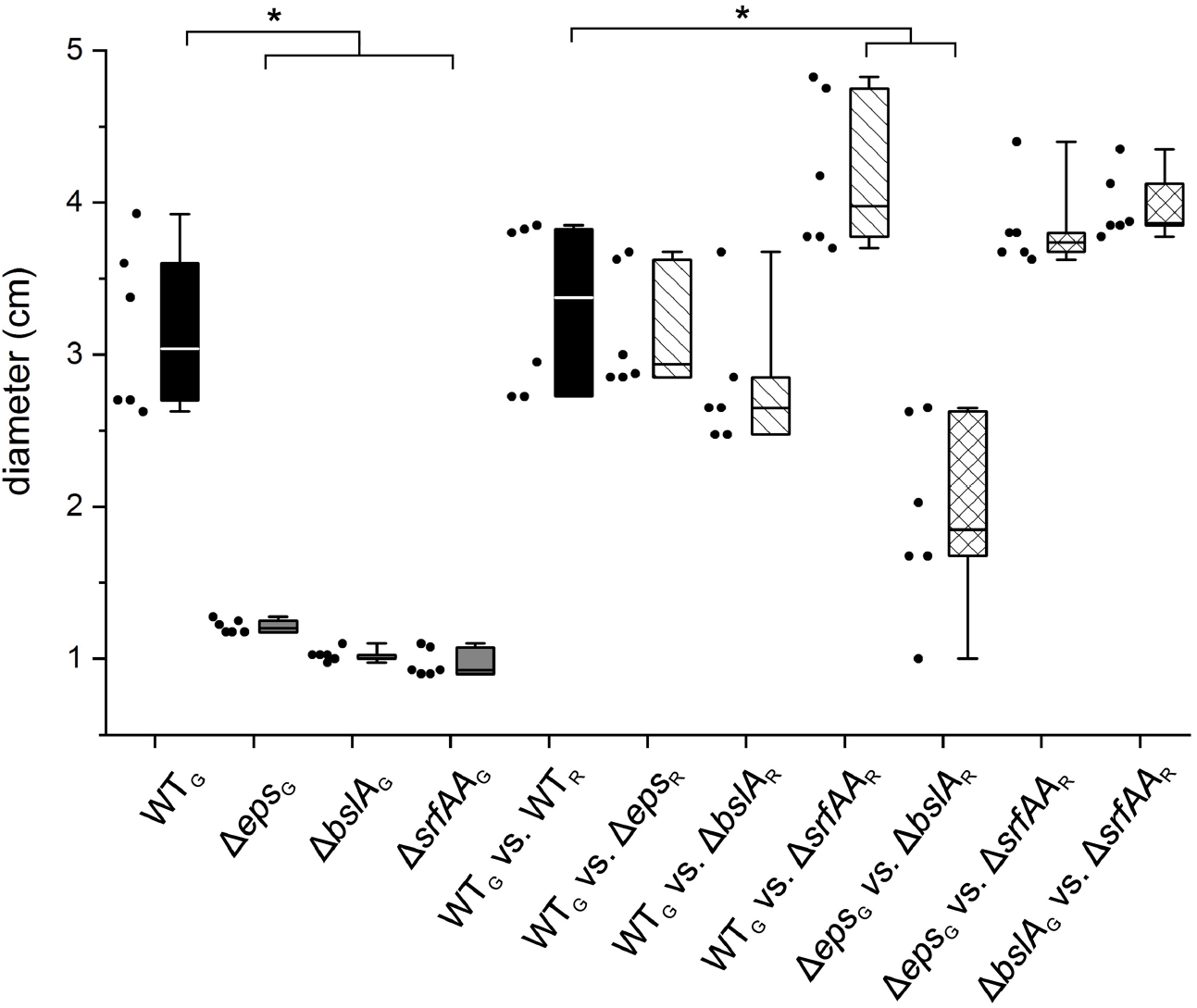
Competition of sliding good producers and non-producers affects sliding expansion. Diameter of sliding colonies were recorded after incubation on semi-solid medium at 37°C for 24 h. For competition assays, fluorescently labeled wild type (WT) and EPS, BslA and surfactin non-producers (Δ*eps*, Δ*bslA*, Δ*srfAA*, respectively) were mixed at 1:1 initial ratio (G – green fluorescent strain, R – red fluorescent strain). The box indicates the 25th-75th percentile; the line in the boxes represents the median. Single data values are represented as dots. Asterisks indicate a significant difference to the WT or WT control competition (unpaired two-sample t-test with Welch Correction, P < 0.05, n = 6).

Next, we performed pairwise competition experiments, by first suspending strains in a 1:1 ratio in a liquid culture before inoculating them on the semi-solid agar surface (qualitatively similar results are obtained when mixing 1:10 or 10:1, see Fig S1 and S2; and Supplementary results). To distinguish strains on the surface, we labeled them with distinct constitutively-expressed fluorescent reporters (either GFP or mKate2) and used stereomicroscopy to monitor their distribution within the colony. Swapping the fluorescent reporters between strains had no effect on the outcome of competition (Fig. S3). We first examined how two wild-type strains, only differing in their fluorescent reporters, distribute themselves during colony growth. Strains strongly segregated in space, giving rise to more-or-less evenly distributed sectors growing out from the colony center, through the passive process of sliding motility (Fig. 2A; see also [34]). Since the wild-type strains are identical, besides their reporter gene, sliding motility was unaffected and the colony reached the same size as that of the wild types in isolation.

**Fig. 2.**
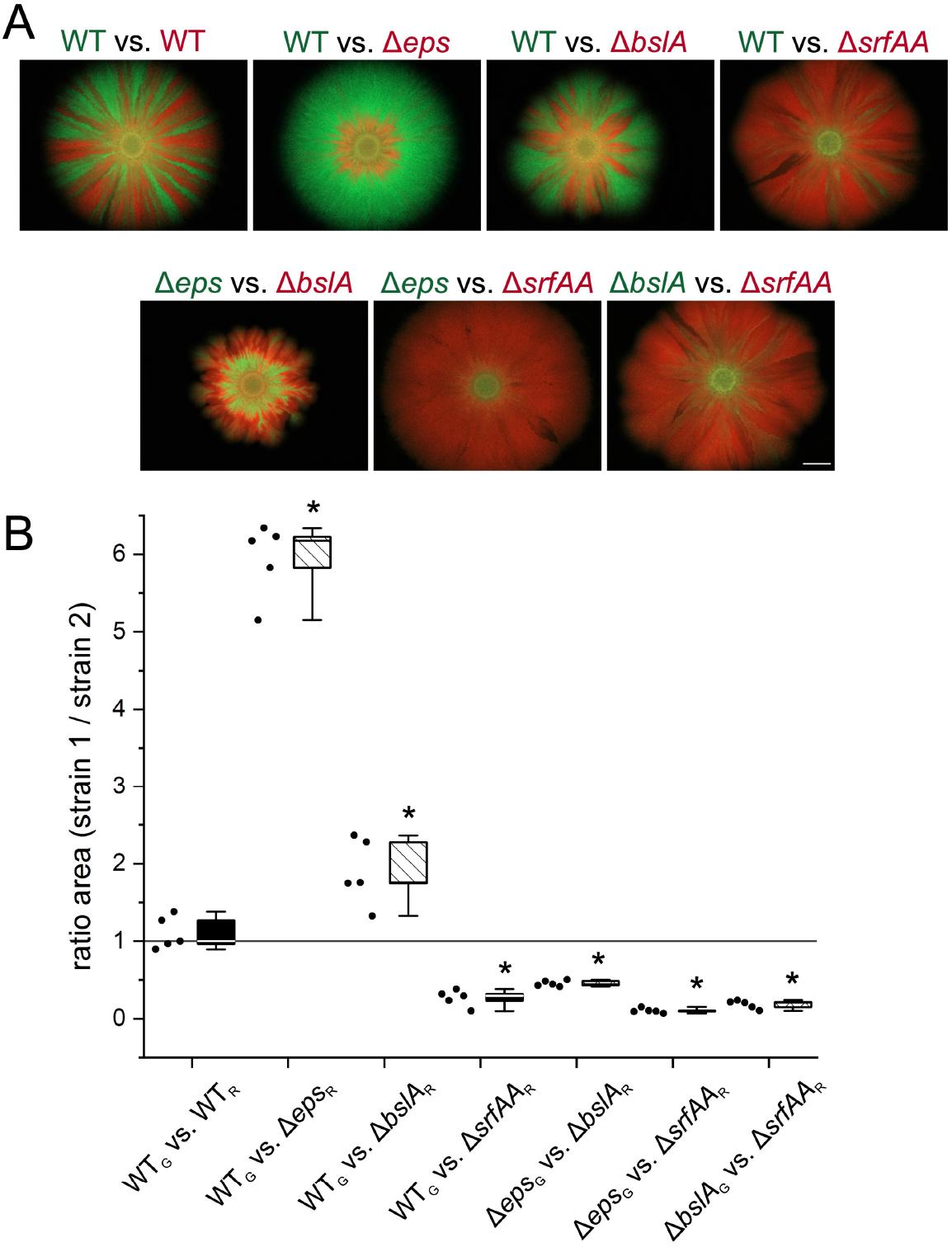
Structure and success of individual strains in competition assay sliding colonies. (A) Overlay of green and red fluorescent images from representative sliding colonies of competition assay from Fig. 1, with 1:1 initial ratio. Green text indicates a green fluorescent strain; red text indicates a red fluorescent strain. The scale is equal for all images and the scale bar represents 5 mm. (B) Ratio of occupied area of strain 1 versus strain 2 (in pixel^2^) of the sliding colonies from (A) obtained by quantitative image analysis using ImageJ. G indicates a green fluorescent strain; R indicates a red fluorescent strain. The box indicates the 25th-75th percentile; the line in the boxes represents the median. Single data values are represented as dots. Asterisks indicate a significant difference to an area ratio of 1 which is marked by a dark grey line (one-sample t-test, P < 0.05, n = 5).

Next, we competed the wild type against each of the non-producing strains. When competing the wild type against the EPS or BslA non-producers, colony expansion was indistinguishable from that of the wild type, i.e. after 24 hours colonies were approximately 3 cm in diameter (Fig. 1). However, in contrast to the wild type, strains did not form radial sectors (Fig. 2A). The non-producing strains were instead confined to the center of the colony and entirely displaced by the wild type at the colony edge. When comparing the surface occupancy of the wild type with either that of the EPS or BslA non-producer, the wild type was 6 and 2 times more abundant, respectively (Fig. 2B, one-sample t-test, P_WT/Δ*eps*_ = 2·10^−5^, P_WT/ΔbslA_ = 0.01, n = 6). This suggests that the EPS and BslA non-producer cells cannot exploit the wild type in competition for space. In contrast, the surfactin non-producer does seem to exploit the wild type: in competition with a surfactin non-producer, the wild type was largely displaced at the colony edge and the surface occupancy was about three times smaller than that of the surfactin non-producer (Fig. 2, one-sample t-test, P < 0.05, n = 5). Interestingly, colonies consisting of both wild type and surfactin non-producers were about 1 cm larger in diameter than wild-type colonies (Fig. 1, unpaired two-sample t-test with Welch Correction: P = 0.043, n = 5). Since sliding motility is driven by cell division, these results suggest that surfactin non-producers have a significant growth advantage over the wild type and can advance colony expansion by exploiting surfactin production by the wild type.

Finally, we also competed each of the non-producers against each other. In all cases, colony growth was either partly or completely recovered when non-producers were grown together in comparison to their growth in isolation (Fig. 1). This suggests that to a certain degree all secreted molecules are being shared. Complete recovery of impaired growth was only observed when Δ*srfAA* was cultured with either Δ*eps* or Δ*bslA*, and partial recovery was observed for the co-culture of Δ*eps* and Δ*bslA*. Like for the wild type, the surfactin non-producer outcompeted both the EPS and BslA non-producers (Fig. 2, one-sample t-test, P < 0.05, n = 5), further corroborating the idea that surfactin acts like a common good that can be exploited. When competing Δ*eps* against Δ*bslA*, the BslA non-producer had a competitive advantage, occupying about twice as much space as the EPS non-producer after 24 hours (Fig. 2, one-sample t-test, P < 0.05, n = 5). This suggests that BslA is at least partly shared, enabling Δ*bslA* to exploit Δ*eps*, even though it was unable to exploit the wild type.

Given that sliding is at least partly recovered for all pairwise combinations of non-producers, we conclude that surfactin and BslA can be shared between cells within the colony and EPS potentially as well. Yet, from the three secreted molecules, only surfactin fits the description of a typical common good: Δ*srfAA* cannot slide by itself, but in combination with either the wild type or the other non-producers sliding is fully recovered with Δ*srfAA* having a competitive advantage. In contrast, both Δ*eps* and Δ*bslA* do not have a competitive advantage in competition with the wild type. To further disentangle what determines the outcome of competition, we next quantified the costs of molecule synthesis and assessed the degree by which molecules are shared.

### Cost of secreted molecule production

Since the complex spatial structure of colonies withholds us from directly measuring the costs of molecule production during sliding motility, we measured these costs indirectly by artificially inducing molecule production and examining the costs under liquid growth (using the same growth medium as for the sliding experiments). To this end, we first created strains with an IPTG-inducible promoter in front of the gene or operon responsible for production of the respective extracellular molecules (see Materials and Methods). Using these inducible gene constructs, we then determined what induction level best mimics molecule production in the wild type. We did so by comparing sliding motility under a range of induction levels and determining what induction level best mimics sliding motility in the wild type – this is different for each of the three molecules (Fig. S4). Finally, we measured the production cost by directly competing a non-producing knockout mutant against the inducible strain with and without induction and, then, measuring the relative fitness of the inducible strain relative to the knockout (Fig. 3). Strikingly, only induced surfactin production (+IPTG) was associated with a significant decrease in growth rate (i.e. relative fitness), indicating that in our medium conditions only surfactin production bares significant energetic costs (Fig. 4). These costs lead to a competitive disadvantage and explain why Δ*srfAA* has a competitive advantage over surfactin-producing strains during sliding. In contrast, and to our surprise, we did not observe any measurable costs for EPS and BslA production under sliding conditions, suggesting that these molecules are cost-free and cannot be exploited.

**Fig. 3.**
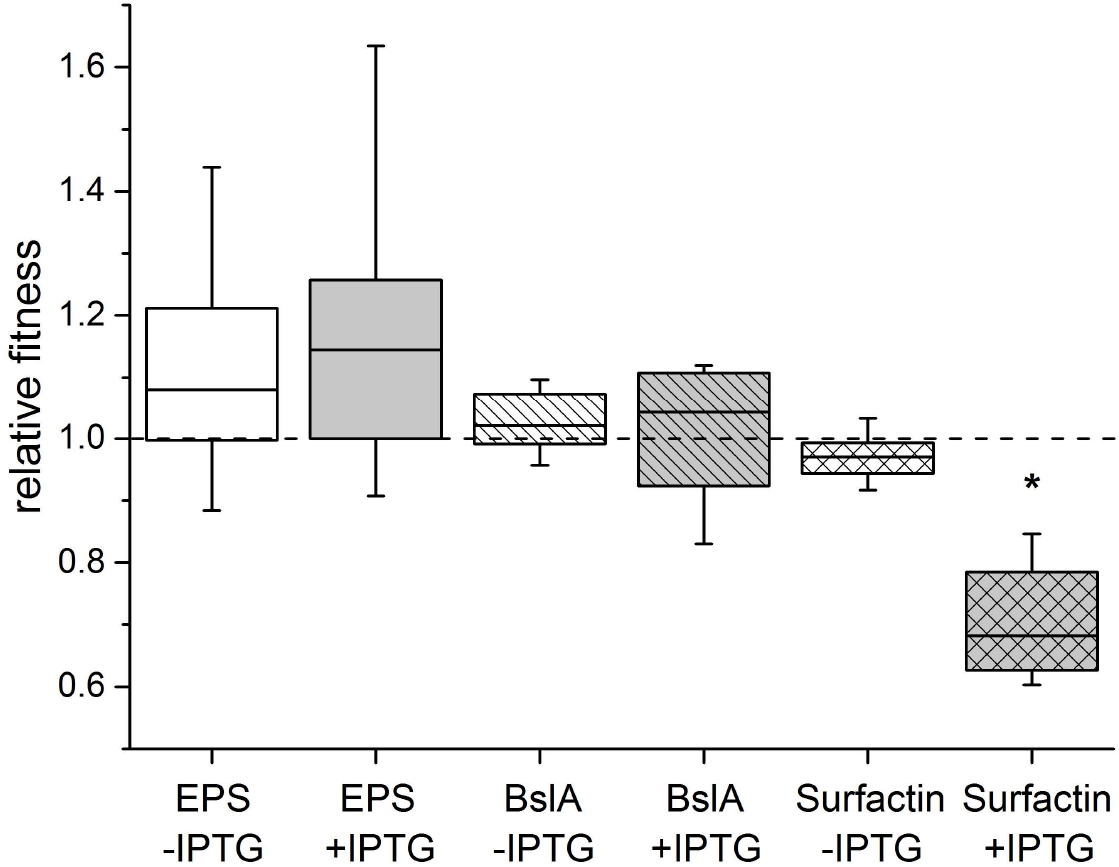
Surfactin is costly to produce under sliding promoting conditions. Relative fitness calculated from initial and final cfu of competitions between a good non-producer against the respective IPTG-inducible strain. The competition was conducted in IPTG containing medium (+IPTG) and as control in medium lacking IPTG (-IPTG). The box indicates the 25^th^-75^th^ percentile; the line in the boxes represents the median. The dashed line indicates a relative fitness of 1. The asterisk indicates a significant difference to 1 (one-sample t-test, test mean = 1, P = 6.9·10^−4^, n = 6).

**Fig. 4.**
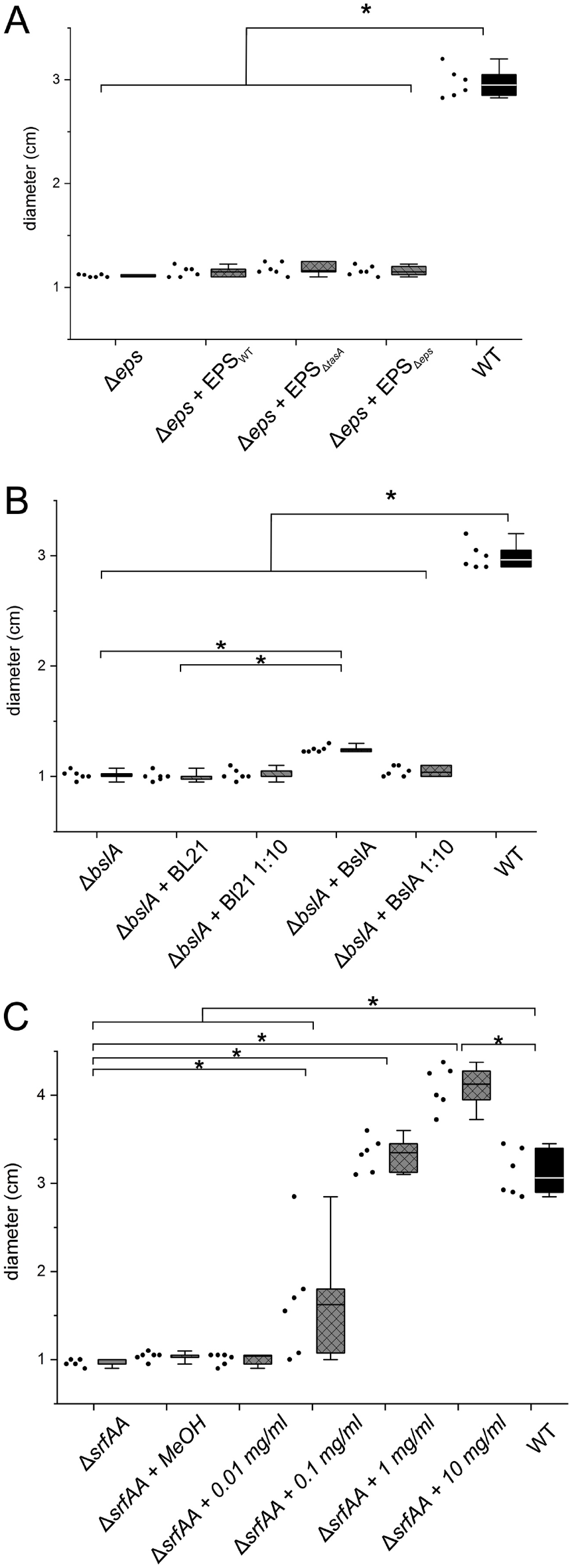
Complementation of non-producer sliding with supplied molecules. Diameter of EPS (A), BslA (B) and surfactin (C) non-producers (Δ*eps*, Δ*bslA*, Δ*srfAA* respectively) incubated under sliding promoting conditions with supplied molecules or a control substance and wild type controls for 24 h. For EPS non-producer complementation (A), EPS isolated from wild type, *tasA* mutant or *eps* mutant biofilms was used. For BslA non-producer complementation (B), BslA containing supernatant from an *E. coli* strain harboring a BslA production plasmid was used (+BslA); supernatant from an *E. coli* strain without the plasmid served as control (+BL21). For surfactin non-producer complementation (C), commercially available surfactin was used in different concentrations (0.01, 0.1, 1, and 10 mg/ml) dissolved in methanol, which was used as control (+MeOH). The box indicates the 25^th^-75^th^ percentile; the line in the boxes represents the median. Single data values are represented as dots. Asterisks indicate significant differences.

### Complementation by the secreted molecules

Since colony expansion was (partly) recovered when mixing different non-producing mutants (Fig. 1 and 2), we know that secreted molecules are at least partially shared. Next, we investigated to which extend they are shared, by directly adding EPS, BslA or surfactin to colonies of Δ*eps*, Δ*bslA* and Δ*srfAA* mutants and examining if and to what degree colony expansion was recovered. We isolated EPS, BslA and surfactin from different sources. For EPS, we extracted EPS-containing matrix from either wild-type biofilms or Δ*tasA* biofilms; these latter biofilms lack the main protein component in the biofilm matrix (i.e. TasA). As a control, we also collected matrix from Δ*eps* biofilms, which lacks EPS. BslA was isolated from the supernatant of a *E. coli* BL21 strain that was engineered to overproduce BslA. The quality and functionality of our extraction methods for both EPS and BslA were validated experimentally (see Fig. S5, S6). For surfactin, we simply used commercially available purified surfactin (Sigma Aldrich; CAS# 24730-31-2).

We next supplemented colonies of non-producing mutants with different concentrations of external EPS, BslA and surfactin and monitored their growth for 24h. Δ*eps* colonies did not show any recovery of colony expansion when adding EPS, while Δ*bslA* colonies showed a slight increase in size when adding BslA (Fig. 4A, P > 0.05 and Fig. 4B, P_mutant_ = 1.2·10^−6^, P_BL21_ = 9.4·10^−7^, P_WT_ = 7.6·10^−8^, respectively; all unpaired two-sample t-test with Welch Correction, n = 6). However, this effect was strongly dependent on the BslA concentration: when adding 10-fold diluted BslA-containing supernatant no increase in colony size was observed (unpaired two-sample t-test with Welch Correction, P_BslA1:10_ = 0.21, n = 6). In contrast, when adding surfactin to Δ*srfAA* colonies, we observed full recovery of colony expansion; at high surfactin concentrations colony expansion of Δ*srfAA* colonies even exceeded that of the wild type (Fig. 4C, P_1 mg/ml_ = 0.049, P_10 mg/ml_ = 6·10^−5^, P_WT-10 mg/ml_ = 6·10^−5^; all unpaired two-sample t-test with Welch Correction, n = 6).

These results are consistent with our competition experiments: surfactin acts like a common good that is freely shared within the colony, while EPS and BslA are not or only marginally shared and therefore act as private goods that mostly benefit the producer. As a consequence, the wild type outcompetes both Δ*eps* and Δ*bslA* mutants during surface competition. At the same time, we do observe that Δ*bslA* mutant can partly be compensated by externally added BslA, which explains why Δ*bslA* somewhat profits from growing together with Δ*eps* (Fig. 2), leading to a partial recovery of colony expansion.

### Agent-based modeling recapitulates the observed complementation dynamic

Since we are unable to manipulate diffusion rates (i.e. degree of sharing) and private benefits of EPS and BslA production experimentally, we decided to further investigate how these factors impact sliding motility using an agent-based model. Following our experiments, we model surface competition during sliding motility, where colony expansion is driven by cell division. The colony can only expand when the concentrations of surfactin, EPS and BslA are sufficiently high. Wild type cells secrete all three molecules at a constant rate, and incur a small fitness penalty (∼3%) for surfactin production (relative to a Δ*srfAA* mutant; like quantified experimentally). Molecules also degrade at a constant rate and diffuse in space. The molecule concentration determines whether (locally) the colony can expand (see Materials and Methods). Finally, we also assume that EPS and BslA producers can convey private benefits to producers, by increasing their propensity to expand. We model this by lowering the EPS and BslA concentrations at which EPS and BslA producers expand outwards. In our model, we manipulate both the degree by which molecules are being shared, i.e. diffusion rates (10^−8^, 10^−6^, 10^−4^ or 10^−2^ mm^2^ s^-1^), and their private benefits (0%, 10%, 50% or 90%), thereby determining how these factors affect the outcome of surface competition. For each parameter setting, we performed all pairwise competition experiments (Fig. 2): (1) WT vs. WT, (2) WT vs. Δ*eps*, (3) WT vs. Δ*bslA*, (4) WT vs. Δ*srfAA*, (5) Δ*eps* vs. Δ*bslA*, (6) Δ*eps* vs. Δ*srf*, (7) Δ*bslA* vs. Δ*srf*. We then scored the accuracy of our model predictions, both in terms of colony sizes and fitness values (see Materials and Methods): 0 means that none of the pairwise competition experiments were correctly predicted by our model; 7 means that all of them were correctly predicted.

Although diffusion affects competition, Fig. 5 shows that private benefits of EPS and BslA production were most important in predicting the outcome of competition. The best match between our simulation and experimental data (see asterisk in Fig. 5) occurs for relatively high private benefits of EPS production (∼90%) and relatively low but nonzero private benefits for BslA production (∼10%). Similarly in agreement with our data, we find considerable variation between replicate simulations (e.g. Fig. 1), which likely result from so-called founder effects that are commonly observed in (bacterial) surface expansion [39, 40]. Fig. 6 shows the simulation results that most accurately match our experimental results (asterisk in Fig. 5). In accordance with our experimental findings, we observe that EPS mostly accumulates locally and therefore barely shared, while BslA shows limited diffusion and surfactin diffuses freely within and beyond the colony. The distribution of strains after 24h of colony growth closely matches our experimental findings (Fig. 2).

**Fig. 5.**
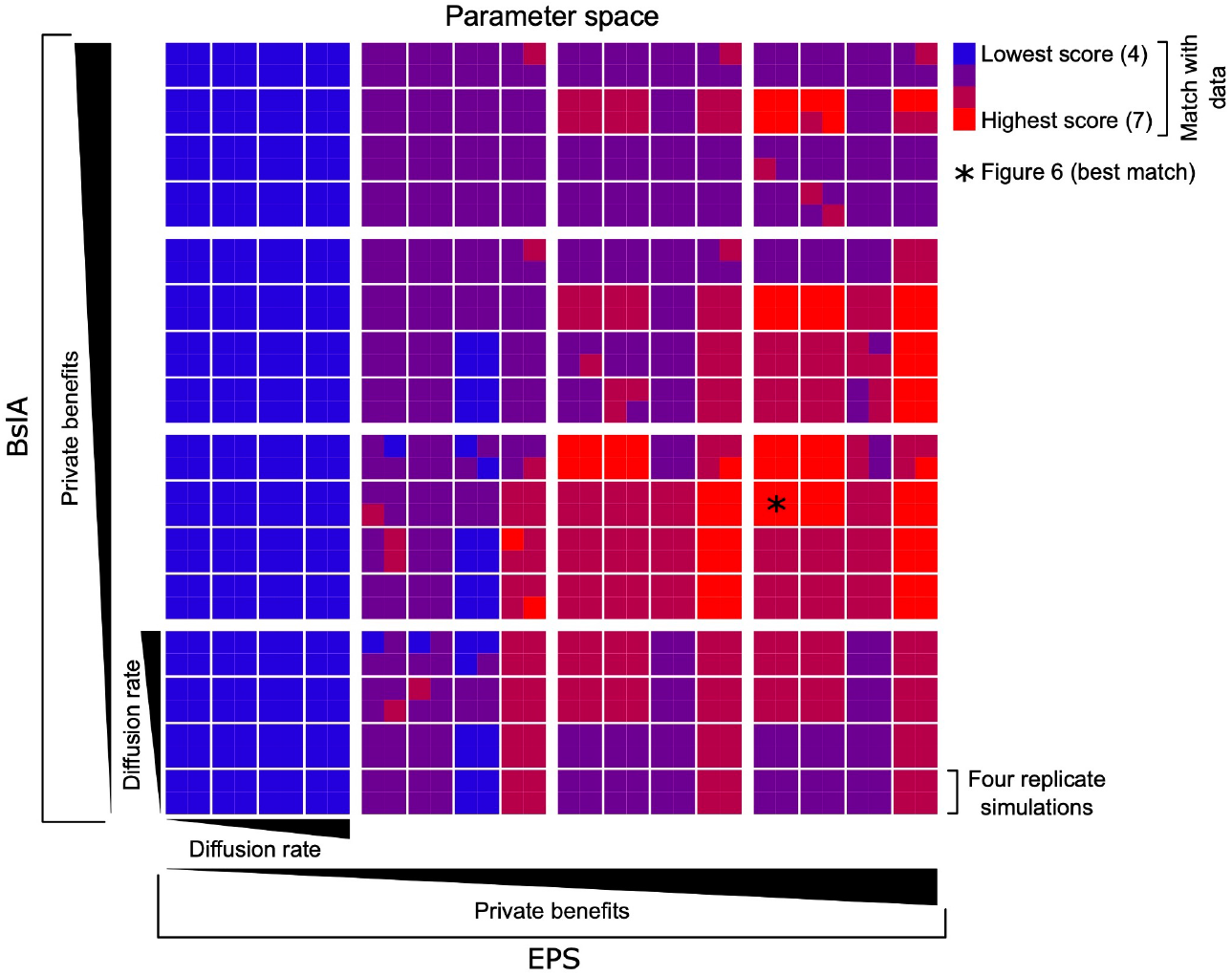
Effects of private benefits and diffusion rates of EPS and BslA on outcome of competition. Smallest blocks show four attached squares (2×2), which represent replicate simulations; intermediate block show 4×4 smaller blocks that differ in diffusion rates; largest blocks contain 4×4 intermediate blocks that differ in private benefits. Asterisk shows what parameter conditions lead to model results that best recapitulate those of the experiments. For model description and parameters see Methods and Supplemental material.

**Fig. 6.**
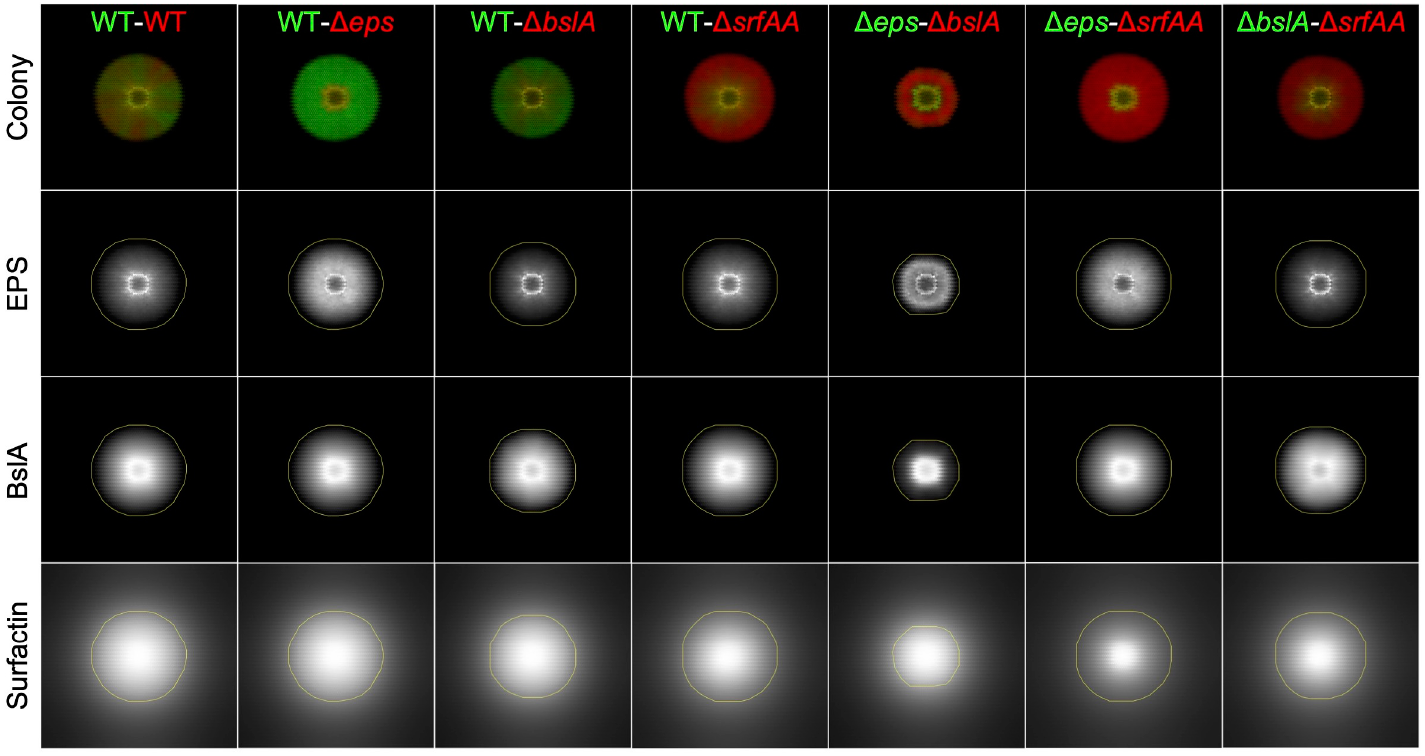
Modeling results of sliding motility. Upper panel, outcome of pairwise competition between wild type (WT), surfactin non-producer (Δ*srfAA*), EPS non-producer (Δ*eps*) and BslA non-producer (Δ*bslA*). Lower panels, concentration of EPS, BslA and surfactin, normalized between the 0 (black) and highest (white) simulated concentrations.

## Discussion

Surface competition plays a critical role in the ecology of bacteria. Accordingly, bacteria often secrete molecules that facilitate surface spreading, enabling them to monopolize space in competition for resources with other bacteria that co-occur on the surface. *B. subtilis* secretes EPS, BslA and surfactin to facilitate sliding motility, a rapid mode of surface spreading that is driven by cell division. Here, we reveal the complex extracellular biology of sliding motility, where EPS, BslA and surfactin have a vastly different impact on surface competition. Although each of these molecules is secreted by cells and essential for sliding motility, only surfactin is costly to produce and diffuses broadly through the colony – thereby acting like a common good. The other two types of molecules are only partly shared, cost-free and provide direct benefits to the producer. These distinct functional consequences of molecule production impact both bacterial ecology and evolution.

Since surfactin acts like a common good, the wild type is outcompeted by Δ*srfAA* during surface competition. Interestingly, we also observed that chimeric WT-Δ*srfAA* colonies show more efficient sliding motility than wild-type colonies, which likely results from the fact that Δ*srfAA* mutants divide quicker than the wild type and thereby accelerates colony expansion. Thus, the presence of cells that do *not* produce surfactin can benefit overall colony expansion. Similarly, MaClean *et al*. [41] previously demonstrated that, in structured yeast populations, productivity during growth on sucrose was maximized when mixing cells that produce an enzymatic common good (invertase) with those that do not. Interesting, even in wild-type *B. subtilis* colonies, not all cells produce surfactin – due to heterogeneous gene expression – suggesting that even these colonies partly mitigate the costs of surfactin production through a phenotypic division of labor [8, 26, 42]. A genetic division of labor between wild-type and Δ*srfAA* cells in chimeric colonies lies in extension of this [24]. It might therefore not be surprising that surfactin mutants are frequently isolated from soil samples that also contain surfactin producers [43]. It is however yet unclear how long wild type and surfactin mutants can stably co-existence in time (i.e. without displacement of the wildtype).

If surfactin production can so easily be exploited, what could prevent wild-type cells from being outcompete by surfactin mutants in nature? Here, transcriptional regulation of the surfactin genes probably plays a crucial role: their expression is controlled through quorum sensing, using the strain-specific pherotype ComX [44, 45]. Quorum-sensing regulation has two possible effects. First, it prevents wasteful production of surfactin at low cell densities – when there is no benefit of producing surfactin. Second, surfactin could specifically be expressed when cells are surrounded by kin, similar or the same strain, which reduces the risk of exploitation [46, 47]. Finally, other ecological factors could play a role in preventing surfactin producers from being outcompeted. For example, besides facilitating sliding motility, surfactin also acts like an antimicrobial that kills other bacterial species. Surfactin producers might therefore gain an ecological advantage in competition with other species [48].

Although both EPS and BslA are secreted [15, 16, 27, 37], both the experiments and model suggest that these molecules are not broadly shared and provide direct private benefits to the producers. Consistently, a previous study of van Gestel and colleagues [6] showed that *eps* knockout mutants are excluded from the EPS-producing filamentous bundles at the colony edge during sliding motility. In contrast, under biofilm or pellicle growth conditions, EPS was previously shown to act as a common good that is broadly shared [22, 24, 26, 49, 50]. This striking difference in the functional consequences of EPS production during sliding motility and complex colony formation (e.g. biofilm and pellicle formation) shows that one should be cautious in qualifying molecules as generic common goods: molecules can act as classical common goods in one condition and as private goods in another. Thus, rather than determining whether molecules act as common goods, the future challenge lies in understanding what factors determine the costs, shareability and private benefits of molecules in different ecological settings.

There are several factors that we did not study that could further influence the outcome of surface competition. For example, secreted molecules might alter gene expression and therefore lead to indirect phenotypic changes that impact competitive fitness. Zhang and colleagues demonstrated that matrix producing cells have a stronger response to quorum sensing signals, thereby altering their gene expression that could lead to indirect phenotypic benefits [51]. Moreover, cells could directly influence the diffusion of secreted molecules and manipulate their private benefits. For example, siderophores secreted by *Pseudomonas aeruginosa* were shown to bind to amyloid fibrils, thereby limiting their diffusion and hence exploitation by non-producers [52]. By the same token, Psl exopolysaccharide secreted by *P. aeruginosa* anchor to the cell in early stages of biofilm formation [53]. Similar retention mechanisms might also play a role in the privatization of EPS during sliding motility.

In many ways, we are only at the beginning of exploring the extracellular biology of bacteria. Advances in single-cell microscopy, transcriptomics and molecule tracking will likely continue to push our understanding of surface-bound microbial communities [21, 54, 55]. In addition, we believe important challenges lie in developing laboratory settings that most accurately reflect the complex ecologies to which bacteria are exposed in nature [56].

## Acknowledgements

We thank Nicola Stanley-Wall for providing strain NRS4110. We thank Christian Kost for his valuable suggestions during the project. This work was funded by the Deutsche Forschungsgemeinschaft (DFG) to Á.T.K. (KO4741/3.1) and by the Danish National Research Foundation (DNRF137) for the Center for Microbial Secondary Metabolites. T.J. was supported by the International Max Planck Research School “The Exploration of Ecological Interactions with Molecular and Chemical Techniques” and FAZIT Stiftung. J.v.G. was supported by a Marie Sklodowska-Curie Individual Fellowship (742235) and a Swiss National Science Foundation Postdoc Mobility fellowship (P400PB_186789).

