## Supplementary materials for "Complex extracellular biology drives surface competition in *Bacillus subtilis*"

### Supplemental material

**Table S1.** Strains used in this study

| Strain | Genotype | Reference |
|---|---|---|
| *Bacillus subtilis* DK1042 (NCBI 3610 naturally competent derivative) | | |
| TB530 | $P_{hyperspank}$-*gfp:*Spec *hag::*Km | Mhatre *et al.*, 2017 |
| TB531 | $P_{hyperspank}$-*mKATE2:*Spec *hag::*Km | This study |
| TB532 | $P_{hyperspank}$-*gfp:Spec hag::Km eps::Tet* | This study |
| TB533 | $P_{hyperspank}$-*mKATE2:*Spec *hag::*Km *eps::*Tet | This study |
| TB534 | $P_{hyperspank}$-*gfp:*Spec *hag::*Km *bslA::*Cm | This study |
| TB535 | $P_{hyperspank}$-*mKATE2:*Spec *hag::*Km *bslA::*Cm | This study |
| TB536 | $P_{hyperspank}$-*gfp:*Spec *hag::*Km *srfAA::*MLS | This study |
| TB537 | $P_{hyperspank}$-*mKATE2:*Spec *hag::*Km *srfAA::*MLS | This study |
| TB873 | $P_{spachy}$-*bslA:*Spec *bslA::*Cm *hag::*Km | This study |
| TB875 | $P_{hyperspank}$-*epsA-O:*Spec *hag::*Km | This study |
| TB977 | $P_{hyperspank}$-*srfA:*MLS *hag::*Tet | This study |
| TB893 | *hag::*Km *eps::*Tet | This study |
| TB922 | *hag::*Tet *bslA::*Cm | This study |
| TB895 | *hag::*Km *srfAA::*MLS | This study |
| DK1042 | *comI*$^{Q12L}$ | Konkol *et al.*, 2013 |
| TB601 | *eps::*Tet | (Dragoš *et al.*, 2017) |
| TB602 | *tasA::*Km | (Dragoš *et al.*, 2017) |
| *Escherichia coli* | | |
| *E.coli* TB887 | MC1061 pTB887 *(*pTB234; *hxlR:lacI:*Phy-*srfA)* | This study |
| *E.coli* | BL21 | Novagen |
| *E.coli* NRS4110 | BL21 pGEX_TEV_$BslA_{42-182}$::Amp | Hobley *et al.*, 2013 |

**Table S2.** Primers

| **Primer** | **Sequence** (5′-3′) |
|---|---|
| oTH1<br>(Mhatre *et al.*, 2017) | GCATCTAGAGTTGCTCGCGGGTAAATGTG |
| oTH2<br>(Mhatre *et al.*, 2017) | CGAGAATTCATCCAGAAGCCTTGCATATC |
| oTH33 | CTGAAGCTTAAGATTAGGGGAGGTATGAC |
| oTH34 | GGAGCATGCAATGTTCCCGTCACAACATC |
| oTH35 | GCAGGTACCGCAATGTTCCCGTCACAACATC |
| oTH36 | CATCTCGAGGTAAATGTGAGCACTCACAATTC |

**Table S3.** Model parameters.

| **Symbol** | **Parameter** | **Value** |
|---|---|---|
| T | Duration of the simulation | 24h |
| x | Width hexagonal grid | 100mm |
| ΔT | Discretization of time | 1 $min^{-1}$ |
| Δx | Discretization of space | ~1 $mm^{-1}$ |
| ø | Diameter of inoculum | 10 mm |
| $R_{init}$ | Resource concentration at onset of growth | 100 |
| $d_1$ | Secretion rate of surfactin | 0.02 $min^{-1}$ $cell^{-1}$ |
| $d_2$ | Secretion rate of EPS | 0.002 $min^{-1}$ $cell^{-1}$ |
| $d_3$ | Secretion rate of BslA | 0.002 $min^{-1}$ $cell^{-1}$ |
| δ | Degradation rate of secreted molecules | 0.001 $min^{-1}$ |
| $\alpha_1$ | Diffusion coefficient of surfactin | $10^{-2}$ $mm^2$ $s^{-1}$ |
| $\alpha_2$ | Diffusion coefficient of EPS | [$10^{-8}$, $10^{-6}$, $10^{-4}$, $10^{-2}$] $mm^2$ $s^{-1}$ |
| $\alpha_3$ | Diffusion coefficient of BslA | [$10^{-8}$, $10^{-6}$, $10^{-4}$, $10^{-2}$] $mm^2$ $s^{-1}$ |
| $\alpha_R$ | Diffusion coefficient of resources | $10^{-3}$ $mm^2$ $s^{-1}$ [#] |
| $r_{WT}$ | Division rate wildtype | exp(log(2.0)/45.0)-1 $min^{-1}$ [##] |
| $r_{srf}$ | Division rate *srf* knockout mutant | 1.000622*$r_{WT}$ $min^{-1}$<br>(Equates a ~3% fitness benefit) |
| τ | Threshold for colony expansion | 100 |
| $p_{surfactin}$ | Privatized benefits of surfactin producer | 0% |
| $p_{EPS}$ | Privatized benefits of EPS producer | [0, 10, 50, 90] % |
| $p_{BslA}$ | Privatized benefits of BslA producer | [0, 10, 50, 90] % |

[#] Comparable to the diffusion of glycerol in water at a temperature of 25°C.
[##] Equal to having cell division rate of one doubling in 45 minutes

**Table S4.** Model variables.

| **Symbol** | **Variable** |
|---|---|
| *cell* | Cell |
| *R* | Resource |
| $M_1$ | Secreted molecule 1, surfactin |
| $M_2$ | Secreted molecule 2, EPS |
| $M_3$ | Secreted molecule 3, BslA |

**Pseudocode**

**Start of simulation:**

1. Setup hexagonal surface area of 100x100 grid elements, which together are 100mm in width (~1mm$^2$ resolution).
2. Homogeneously distribute resource across grid with 1000 units ($R_{init}$) per grid element.
3. Inoculate cells within a radius of 5 mm from the surface center (inoculum diameter, ø=10mm), with 100 cells per grid element.

**Run simulation for 24 hours, every minute do the following updates:**

1. **Diffusion of secreted molecules and nutrients**: Diffusion of surfactin, EPS, BslA and resources, as determined using Euler's approximation.
2. **Degradation and secretion of molecules**: Degradation of molecules at a fixed rate, δ, and secretion of molecules by cells at fixed rates: $d_1$=0.02 min$^{-1}$ cell$^{-1}$ for Surfactin, $d_2$=0.002 min$^{-1}$ cell$^{-1}$ for EPS and $d_3$=0.002 min$^{-1}$ cell$^{-1}$ for BslA. Δ*srfAA*, Δ*eps* and Δ*bslA* knockout mutants fail to produce one of the respective molecules.
3. **Cell division and colony expansion (i.e. sliding)**: When there are sufficient resources, cells divide with a fixed rate of $r_{WT}$, corresponding to one cell division every ~45 minutes. As measured in the lab, Δ*srfAA* knockout mutants are assumed to have a slightly higher division rate, $r_{srf}$, corresponding to a 3% fitness benefit. By definition, each cell consumes a single unit of resources per cell division. When there are insufficient resources, cells cannot divide. When the concentration of surfactin, EPS *and* BslA exceeds a pre-specified threshold (τ = 100), daughter cells move to any of the neighboring grid elements – thereby mimicking lateral colony expansion (i.e. sliding) – otherwise daughter cells remain on the same grid element as their mother cell (vertical colony expansion). Private benefits result in a reduced threshold at which cells expand outwards. For simplicity, we model this by manipulating the threshold concentration that is required for lateral colony expansion. For example, when wild type cells are assumed to have a 20% private benefit from EPS production, it can expand at a 20% lower EPS concentration (i.e. τ = 80 units of EPS) compared to an *eps* mutant cell.

**Supplementary methods**

Strain construction

To create fluorescently labelled strains, the gene of the respective fluorescence marker was coupled with a Spectinomycin resistance cassette in plasmid pWK-Sp (Susanna *et al.*, 2007), resulting in plasmids pTB498 (mKATE2) and pTB497 (GFP) as described in Mhatre *et al.* (2017). Similarly, for pTB498, the $P_{hyperspank}$-*mKATE2* fragment was amplified using primers oTH1 and oTH2 (Mhatre *et al.*, 2017) from plasmid phy-mKATE2 (van Gestel *et al.*, 2014) and was after digestion with XbaI and EcoRI ligated into pWK-Sp (Susanna *et al.*, 2007). The plasmid was transformed into *E. coli* MC1061 and verified by sequencing. *B. subtilis* DK1042 was transformed with the plasmids and successful transformation was verified by PCR and fluorescence microscopy. Subsequently, *hag::*Km as well as *eps::*Tet, *bslA::*Cm and *srfAA::*MLS deletion constructs were inserted in each fluorescent strain using gDNA of GP901 (J. Stülke lab collection) as well as DL1032 (López *et al.*, 2009), NRS2097 (Verhamme *et al.*, 2009) and DL107 (López *et al.*, 2009) resulting in strains TB530 and 531, TB532 and 533 TB534 and 535 and TB536 and 537, respectively.

To create TB873, *B. subtilis* was transformed with plasmid pDRyuaB (Kovács and Kuipers, 2011) harboring a copy of the *bslA* gene (formerly *yuaB*) under control of an IPTG inducible promoter. The insertion in *amyE* was confirmed by PCR and subsequently, deletion constructs *bslA::*Cm and *hag::*Km were inserted by transformation with gDNA of NRS2097 (Verhamme *et al.*, 2009) and GP901, respectively. To create TB875, *B. subtilis* DK1042 was transformed with gDNA of NRS1685 (Verhamme *et al.*, 2009) and GP901. To create TB977, the *srfA* gene was PCR amplified using primers oTH33 and oTH34, digested with HindIII and SphI and inserted into pDR111 (D. Rudner). The fragment *Phy-srfA* was then PCR amplified using primers oTH35 and oTH36, digested with XhoI and KpnI, inserted into pTB886 containing the *lacI* gene and the region upstream of *srfA* and transformed into *E. coli* resulting in pTB887. The plasmid pTB887 was then used to transform *B. subtilis*, successful integration was verified by sequencing and subsequent transformation with gDNA of GP902 (Diethmaier *et al.*, 2011) resulted in strain TB977. For TB893, *B. subtilis* DK1042 was transformed subsequently with gDNA of GP901 and DL1032 (López *et al.*, 2009). To create TB922, *B. subtilis* DK1042 was transformed subsequently with gDNA of NRS2097 and GP902 (Diethmaier *et al.*, 2011). For TB895, *B. subtilis* DK1042 was transformed subsequently with gDNA of GP901 and DL107 (López *et al.*, 2009).

Test of BslA containing lysate and isolated EPS for pellicle formation

To test functionality of the BslA containing lysate and the isolated EPS, both compounds were mixed in a 1:4 ratio with concentrated biofilm promoting MSgg medium (1.3x concentrated,

Branda *et al.*, 2001). As controls for BslA and EPS, PBS buffer and deionized water were used, respectively. The compound supplemented medium was inoculated 1:100 with overnight cultures of the respective mutant (Δ*bslA* or Δ*eps*) in a 24- or 48-well plate and a control with only MSgg medium was inoculated with the wildtype (all strains in Δ*hag* background, TB532, TB534 and TB530, respectively). The cultures were incubated for 2-3 d to allow for pellicle formation and imaged using a Axio Zoom V16 stereomicroscope (Carl Zeiss, Jena, Germany). Additionally, in the BslA test the hydrophobicity of the pellicles was determined by depositing a 5 µl water droplet on the pellicle. The test was determined successful since the isolated EPS from wild-type and Δ*tasA* mutant could promote pellicle formation of the Δ*eps* mutant but not EPS isolated from this mutant (negative control) (see Fig. S4). Likewise, the BslA containing lysate promoted wild-type like wrinkle formation and robustness of the *bslA* mutant and restored its hydrophobic properties which was not the case for the control lysate (see Fig. S5).

**Supplementary results**

*Examination how the initial ratio of strains affects competition.*

Each combination of wild type and non-producers was tested with an initial ratio of 1:10 or 10:1 (Fig. S1) and ratios of the area occupied by each strain were determined (Fig. S2). In general, the same trends could be observed as in the assay with 1:1 initial ratio. Interestingly, when the EPS non-producer was incubated together with an initially ten times higher amount of surfactin non-producer, the resulting ratio of the occupied area was only slightly lower than 1 and just on the border of significant difference (Fig. S2, one-sample t-test, test mean = 1, P = 0.048, n = 5). In the fluorescence images of the sliding colony, the surfactin non-producer seemed to be dominating, but a thin layer of EPS non-producer was spread almost over the complete area of the sliding colony (Fig. S1). This effect might be possibly caused by the low amount of EPS non-producer strain and the therefore diminished abundance of surfactin. In the case of the BslA non-producer, an initial high abundance had more impact on the final sliding colony structure, with a final area ratio of around 1, it was not outcompeted any more by the wild type (Fig. S2, one-sample t-test, test mean = 1: P = 0.65, n = 5). However, its high abundance was clearly restricting the wild type since the sliding colony was smaller and exhibited an undulate rim indicating an overall decrease in sliding (Fig. S1). Similarly, in the 10:1 competition of BslA and surfactin non-producers, the resulting ratio was around 1 and thus showed a decreased advantage of the surfactin non-producer compared to the 1:1 assay (Fig. S3, one-sample t-test, test mean = 1, P = 0.47, n = 5; Fig. 2). We suggest that the high abundance of the BslA non-producer dampened the expansion of the surfactin non-producer, since the sliding colony structure showed expansion only in distinct positions where the surfactin non-producer reached the colony rim and was able to diverge from the BslA non-producer abundant sections (Fig. S1).

#### Supplementary Figures

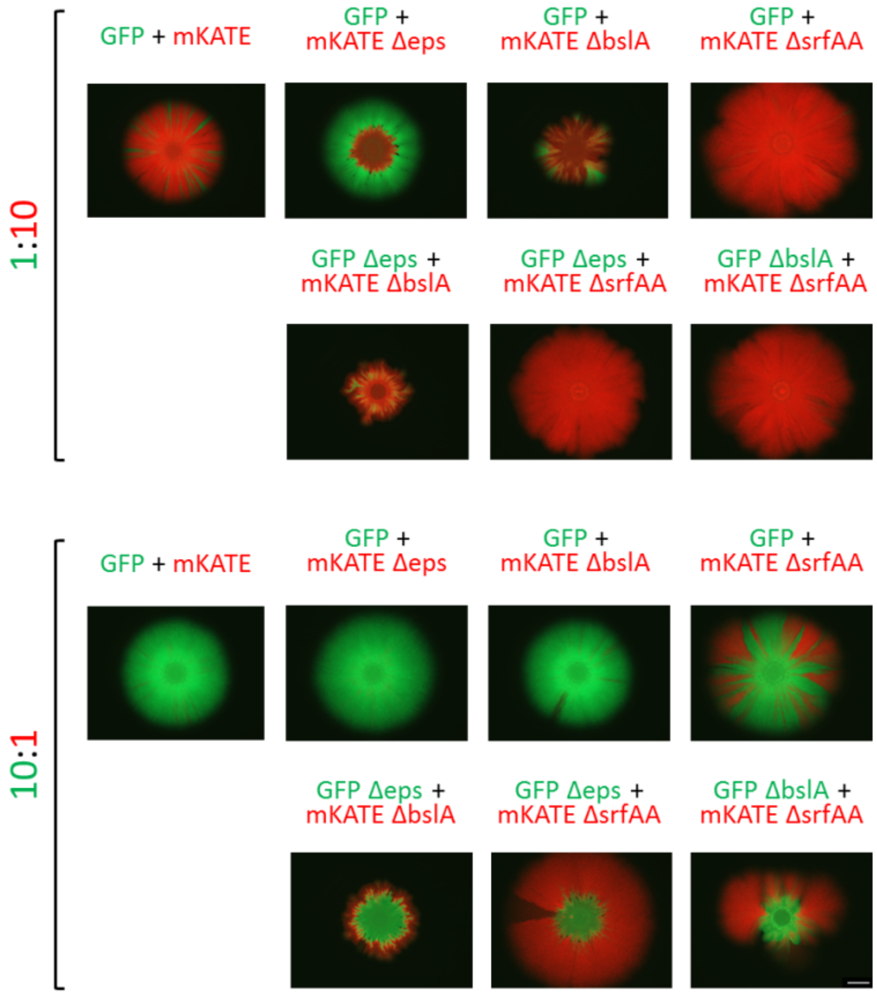


**Fig. S1.** Structure of sliding colonies from competition assays with initial advantage of one strain. Representative overlays of green and red fluorescent images of competition assays with initial ratio of 1:10 (above) and 10:1 (below) are displayed. Green text indicates a green fluorescent strain; red text indicates a red fluorescent strain. The scale bar indicates 5 mm.

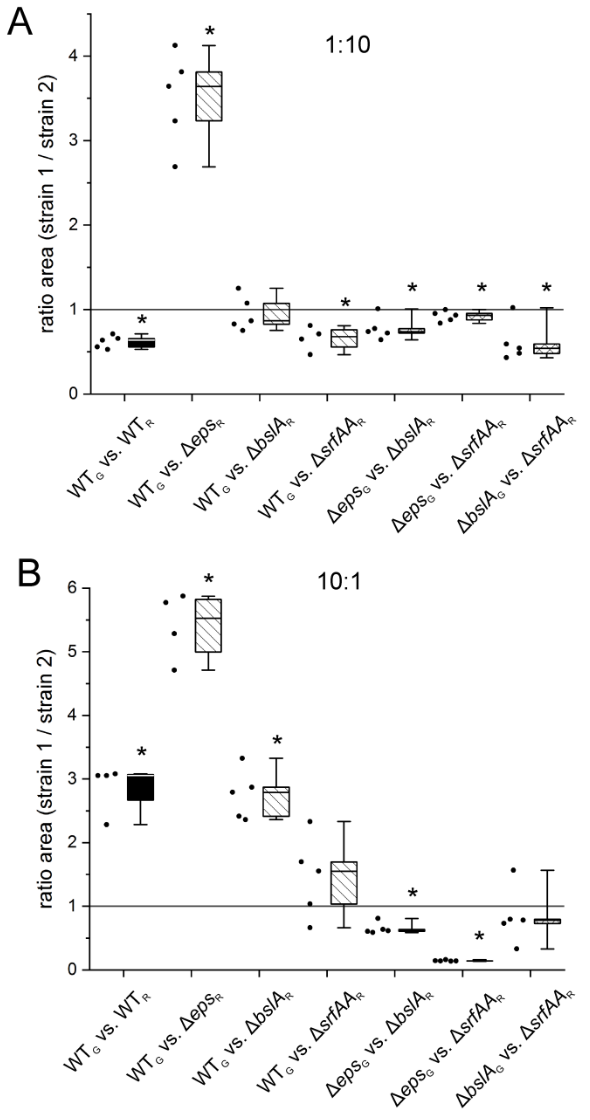


**Fig. S2.** Sliding competition with initial advantage of one strain. Ratio of occupied area of strain 1 versus strain 2 (in pixel$^2$) of sliding colonies from competition assays with 1:10 (A) or 10:1 (B) initial ratio obtained by quantitative image analysis using ImageJ. Strains were incubated on semi-solid medium at 37°C for 24 h. G indicates a green fluorescent strain; R indicates a red fluorescent strain. The box indicates the 25th-75th percentile; the line in the boxes represents the median. Single data values are represented as dots. Asterisks indicate a significant difference to an area ratio of 1 which is marked by a dark grey line.

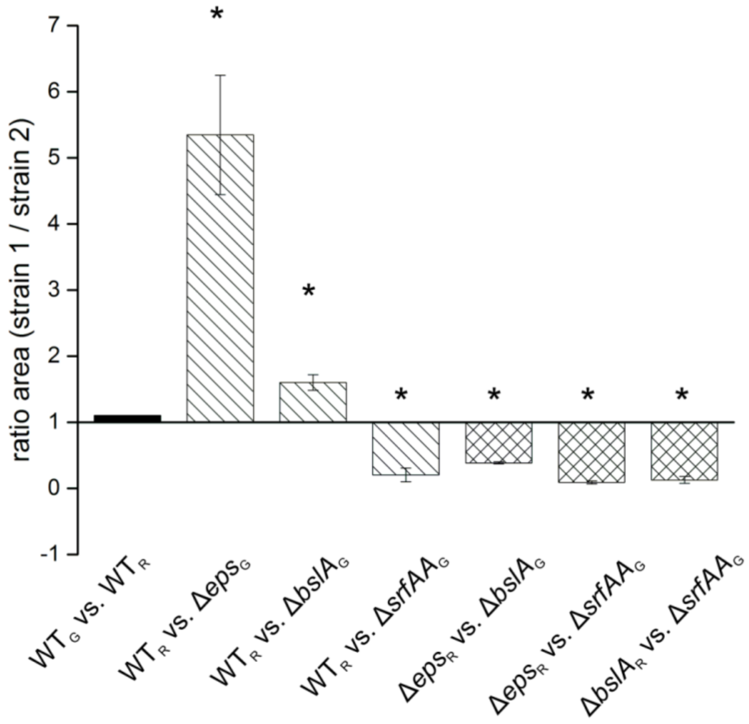


**Fig. S3.** Sliding competition with swapped fluorescence markers. Ratio of occupied area of strain 1 versus strain 2 (in pixel$^2$) of sliding colonies with initial ratios of 1:1 obtained by quantitative image analysis using ImageJ. G indicates a green fluorescent strain; R indicates a red fluorescent strain. Asterisks indicate significant differences to 1 (one-sample t-test, test mean = 1: $P < 0.05$, $n = 5$), error bars indicate the standard deviation.

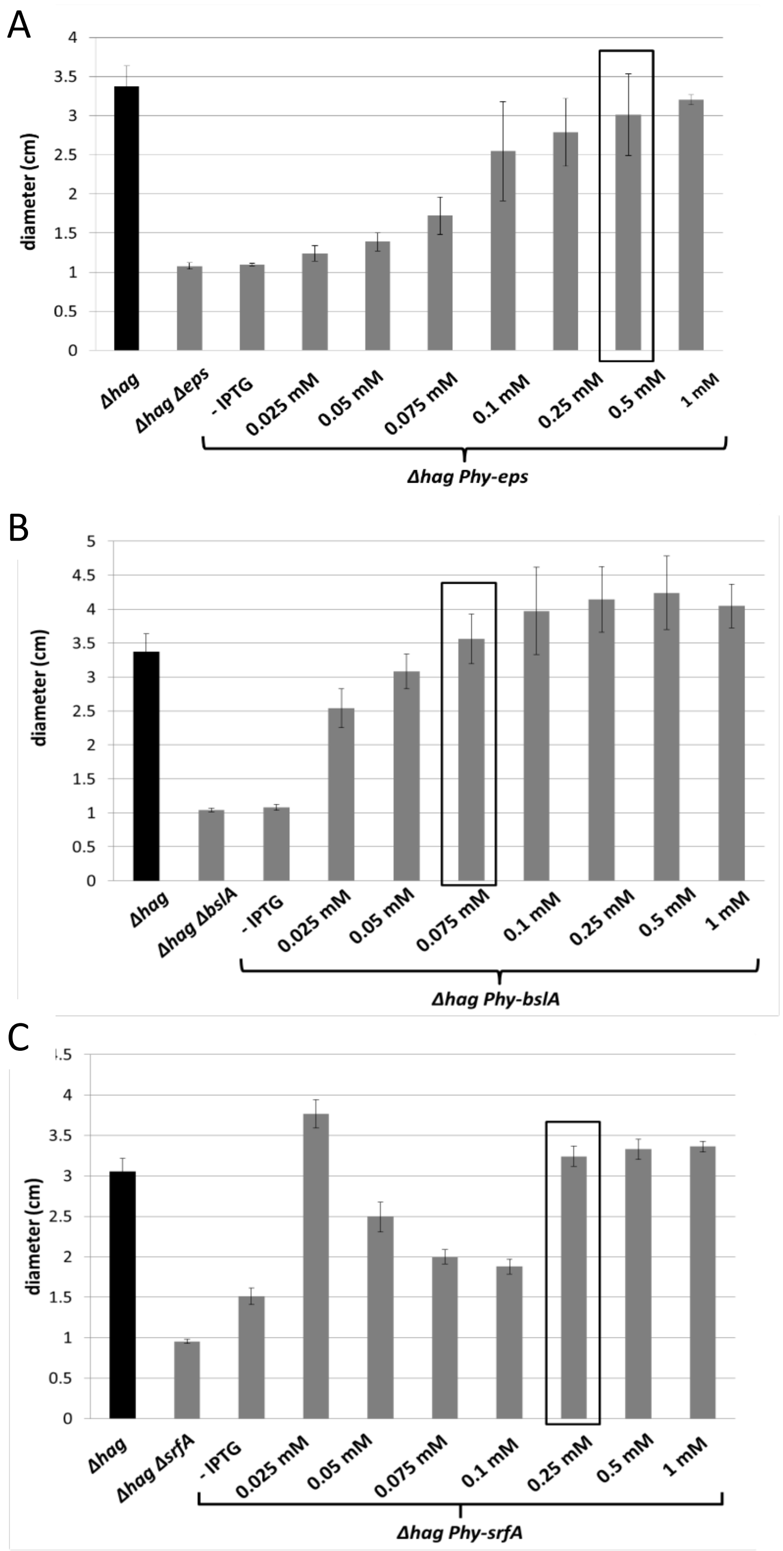


**Fig. S4.** Determination of wild-type level of induction with IPTG. The strains with IPTG inducible (*Phy*) *epsA-O* (A), *bslA* (B) and *srfA-D* (C) genes were tested for wild-type level of sliding at different concentrations of IPTG. The sliding assay was conducted as described in Experimental procedures with IPTG supplemented medium and was evaluated after 24 h by measuring the diameter and was compared to wild-type (Δ*hag*) and the respective mutant. The box indicates the selected IPTG concentration used in the fitness experiments. Note, that the sliding colonies of *Phy-srfA* without and with 0.025-0.075 mM IPTG were partly translucent and showed outgrowth of few denser sectors indicating additional mutations. Error bars indicate the standard deviation (n= 6).

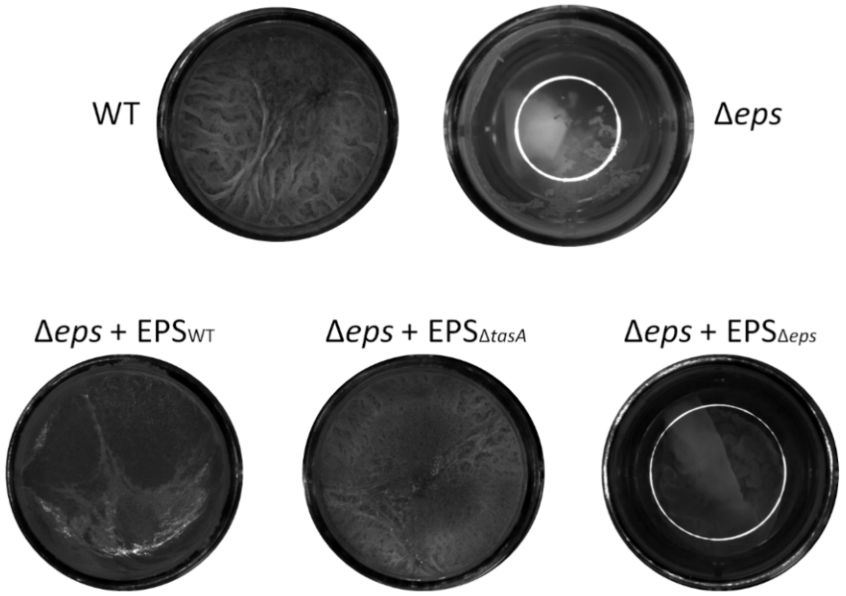


**Fig. S5.** Test of isolated EPS. Strains were incubated statically in concentrated MSgg medium supplemented with deionized water (control WT and Δ*eps*) or isolated EPS from wild-type, Δ*tasA* mutant or Δ*eps* mutant ($EPS_{WT}$, $EPS_{\Delta tasA}$, and $EPS_{\Delta eps}$, respectively) for 2-3 d. The displayed pellicles are representative examples of three or more replicates. Pellicle images were recorded using a Zeiss Axio Zoom stereomicroscope equipped with a black and white camera.

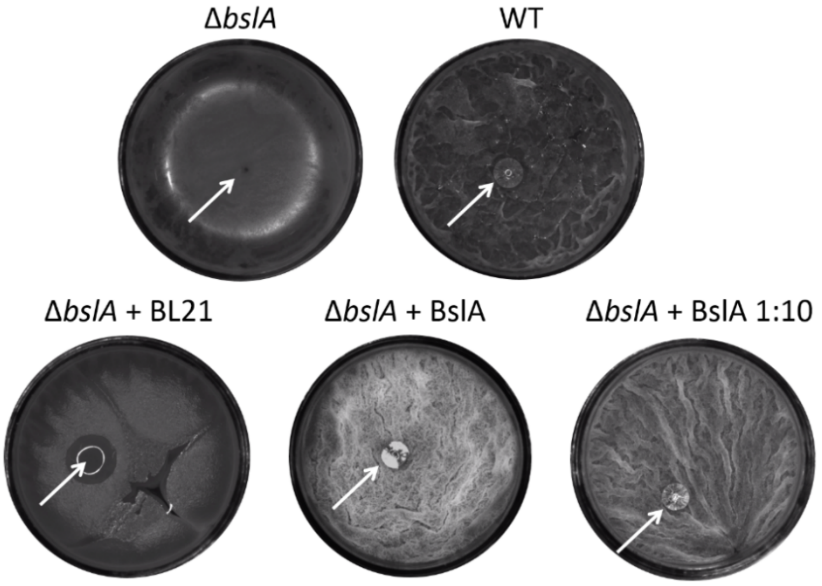


**Fig. S6.** Test of BslA containing lysate. Strains were incubated statically in concentrated MSgg medium supplemented with PBS (control WT and Δ*bslA*), BL21 lysate or BslA containing lysate for 2-3 d. The displayed pellicles are representative examples of three or more replicates and were recorded few seconds after application of a water droplet on the pellicle surface. Images were recorded using a Zeiss Axio Zoom stereomicroscope equipped with a black and white camera and the Camstudio onscreen video program.
